# Mechanobiological proximal tubular defects in ARC syndrome: A *VPS33B* CRISPR knockout study

**DOI:** 10.1101/2025.06.04.657864

**Authors:** Maria Caluianu, Kimberley A. Owen

**Author notes:** Corresponding Author: Maria Caluianu.

## Abstract

ARC syndrome (arthrogryposis-renal dysfunction-cholestasis) is a rare autosomal recessive multisystem disorder affecting the kidneys. The disease is caused by mutations in either *VPS33B* or *VIPAS39*. ARC syndrome is currently incurable, with patients rarely surviving beyond their first year of life. The renal component of this disorder is characterised by proximal tubular dysfunction. Here, a proximal tubular cell line, RPTEC-TERT1, was CRISPR-edited to knock out (KO) *VPS33B* expression. Characterisation of *VPS33B*-KO cells was performed using brightfield imaging, immunostaining, RNA sequencing, and cell detachment assays. The *VPS33B*-KO RPTEC-TERT1 cells demonstrated a ‘peeling’ phenotype and altered cell adhesion. This, along with altered transcription of genes associated with cell adhesion, suggests that *VPS33B* KO results in cell-matrix attachment defects. These findings provide the first insights into the cause of proximal tubular dysfunction in ARC syndrome.

## Introduction

Arthrogryposis–renal dysfunction–cholestasis (ARC) syndrome is a fatal autosomal recessive syndrome caused by a germline mutation in vacuolar protein sorting-associated protein 33B *(VPS33B)* or VPS33B interacting protein, apical-basolateral polarity regulator, spe-39 homolog *(VIPAS39)* (1,2). ARC syndrome is characterised by renal proximal tubular dysfunction, congenital joint contractures, and cholestasis. Currently, there are no treatments for ARC syndrome. Supportive care is offered but most patients die within the first year of life (3).

Approximately 75% of ARC syndrome cases are associated with *VPS33B* mutations (1). Mutations are found throughout the *VPS33B* gene with no evident mutation hotspots (4). The remaining 25% of cases are associated with mutations in the *VIPAS39* gene, again with no evident mutation hotspots (4). These mutations seem to result in a loss of protein function mainly by reducing protein expression and altering the subcellular localisation of the two affected proteins (5–8).

There is limited literature on the renal presentation of ARC syndrome. So far, it is known that patients have proximal tubule dysfunction, with symptoms including aminoaciduria, glucosuria, and renal tubular acidosis (3). Currently, no *in vivo* renal-specific *VPS33B*-knockout (KO) models are available, limiting the study of the renal manifestations of ARC syndrome. Embryonic lethality was reported by E9.5, before the development of the permanent renal metanephros, in global embryonic *Vps33b*-KO mice (9,10). On the other hand, inducible global *Vipas39*-KO or *Vps33b*-KO models induced at adult timepoints failed to demonstrate any visceral abnormalities (11).

Here, we aim to address the lack of model systems in this context and enhance our understanding of the molecular mechanisms underlying the proximal tubular dysfunctions in ARC syndrome by generating and characterising the first *VPS33B*-KO proximal tubular cell line.

## Methods

### Human proximal tubular cell line (RPTEC-TERT1) culture

RPTEC-TERT1s were developed by transfecting normal human adult male primary proximal tubular cells with the human *TERT-1* gene (12). RPTEC-TERT1 cells (ATCC, #CRL-4031™) were cultured in DMEM:F12 Medium (ATCC® 30-2006™), supplemented with hTERT RPTEC Growth Kit (ATCC® ACS-4007™) (1% Supplement A and 1.6% Supplement B), 2% FBS, and 0.1 mg/mL Geneticin™ Selective Antibiotic (G418 Sulfate) (Thermo Fisher, #10131035). Cells were cultured at 37 °C with 5% CO_2_. Media was changed every 2-3 days.

Upon reaching 70-80% confluency RPTEC-TERT1 cells were detached using 0.05% trypsin/EDTA solution, centrifuged at 250 x g for 5 minutes and resuspended at 1:5 density.

### CRISPR-Cas9D10A nickase gene editing in RPTEC-TERT1 cells

RPTEC-TERT1 cells (1.05 x 10^5^ cells/well) were seeded into 6-well plates and transfected with a mixture of 97 µL pre-warmed Opti-MEM, 3 µL FuGENE® 6 Transfection Reagent, and 1 mg/well of either VPS33B Double Nickase Plasmid Set 1 (h1) (Santa Cruz, #sc-406200-NIC) or VPS33B Double Nickase Plasmid Set 2 (h2) (Santa Cruz, #sc-406200-NIC-2) (13). Both plasmid sets contained two plasmids coding for Cas9n (D10A) double nickase, a single guide RNA (sgRNA) sequence (Table 1), and green fluorescent protein (GFP). After 48 hours, GFP-positive, propidium iodide-negative cells were isolated by FACS to identify transfected RPTEC-TERT1 cells with a FACS Vantage flow cytometer (Becton Dickinson Immunocytometry Systems).

**Table 1.**
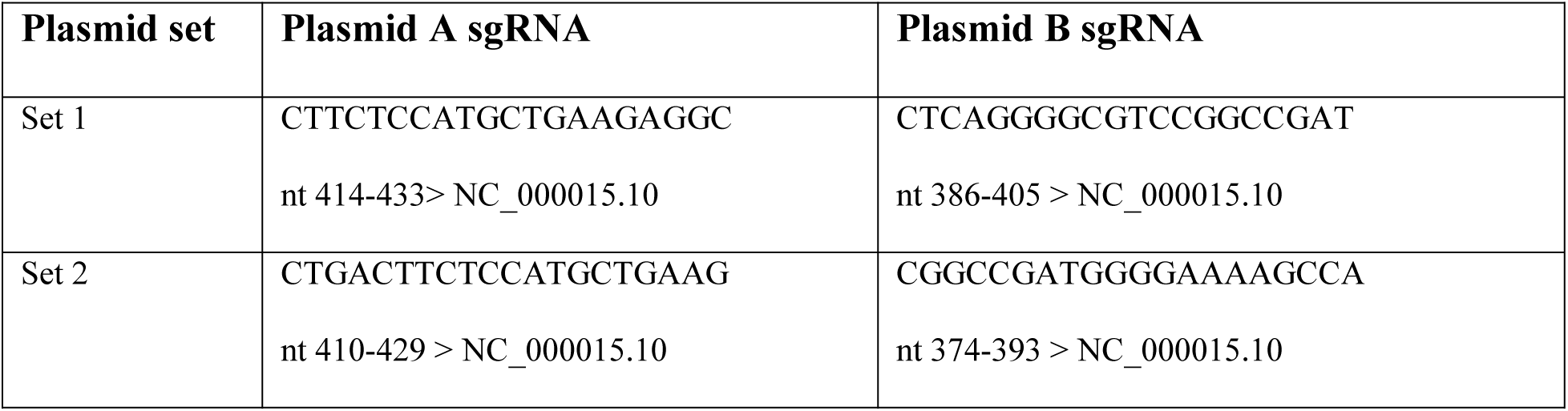
Plasmids used for CRISPR. 20-nucleotide sequences of target-specific sgRNA (single guide RNA) portion of plasmids used for CRISPR knockout of VPS33B (counted from the beginning of the NCBI sequence).

Sorted cells were collected on ice into pre-prepared 15-mL tubes containing 500 µL of fresh medium and transferred to 24-wells, cultured, and expanded. Once the cells had been split into two 6-well plates and reached confluency, media was aspirated from one plate, each well was washed with DPBS and the plate was frozen at -20 °C for DNA extraction for the T7EI assay. The cells from the other plate were passaged into a T25 flask, cultured until 70% confluent, and then single-cell sorted by FACS into individual 96-wells. These cells were cultured for 4-6 weeks without media changes. Once the clones reached confluency, they were split into two new 96-well plates; one for DNA extraction for Sanger sequencing and one for clonal expansion.

### T7 Endonuclease (T7EI) assay

To determine whether a deletion had been created in *VPS33*B, the T7EI assay was used. DNA was extracted from non-transfected and transfected cells, using the GeneJet Genomic DNA Purification kit (Thermo Fisher, #10410450). A 50-µL PCR reaction was then carried out using the human *VPS33B* primers designed to flank the sgRNA cutting sites (Table 2). Then, a 19 µL reaction, composed of 10 µL PCR product, 2 µL 10X NEBuffer™ 2 (Biolabs, #B7002S), and 7 µL MilliQ water, was prepared. The PCR products in this solution were then melted in a thermocycler using the following conditions; 95°C for 5 minutes, ramp down to 85 °C at -2°C/second, ramp down to 25 °C at -0.1 °C/s, and hold at 4°C. 1 µL of T7 Endonuclease I (Biolabs, #M0302S) was added to the reaction and incubated at 37 °C for 1 hour. The T7 cleavage products were then visualised using gel electrophoresis. Successful cleavage of the *VPS33B* gene was indicated by the presence of bands at lower molecular weights than the intact PCR product.

**Table 2.**
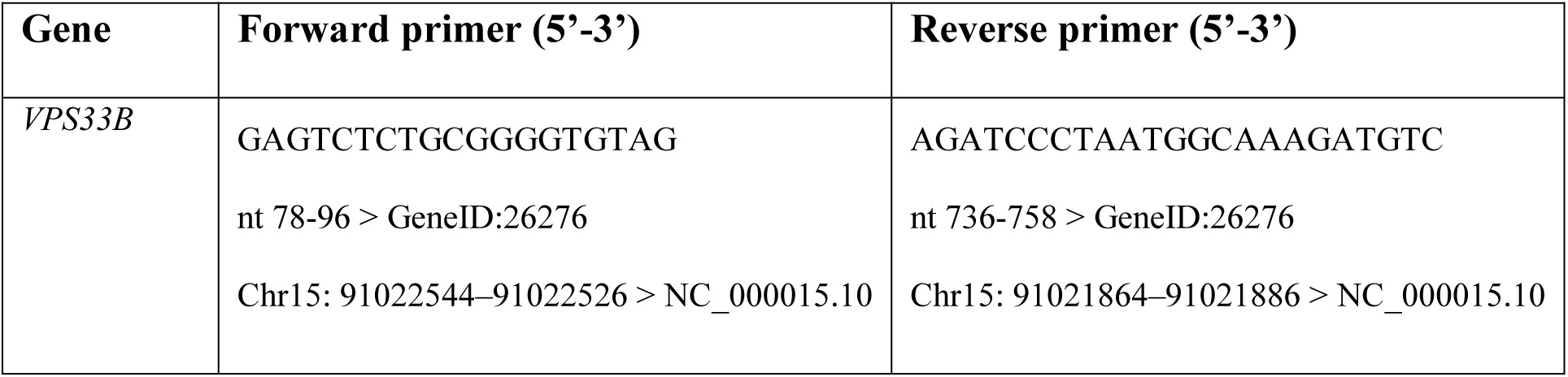
VPS33B primers used for generation of human RPTEC-TERT1 VPS33B-KO CRISPR clones.

### DNA extraction and clonal genotyping

Next, DNA was extracted from the 96-well of CRISPR-transfected clones with QuickExtract™ DNA Extraction Solution (Lucigen, #QE0905T). PCR amplification of the edited region was carried out using the KAPA2G Fast Hot Start and human *VPS33B* primers (Table 2).

### Off-target PCR

To screen for off-target mutations in the transfected clones, PCRs were carried out for four top predicted off-target primer sets. These off-target sites were chosen by screening both sgRNA sequences on COSMID: CRISPR Search with Mismatches, Insertions and/or Deletions setting the maximum number of indels as 2 and allowing for 1 nucleotide insertion or deletion (14) and the WGE CRISPR Finder Tool (15). Potential off-target sites identified by both softwares were selected. Three of these off-target regions were all found in non-coding regions of different chromosomes (Chr20:22233139-22233161, Chr13:82330988-82331010, Chr4:10167787-10167809) while the fourth (Chr1:101898463-101898485) was found to be in the olfactomedin 3 (*OLFM3*) gene. Primer sequences recommended by COSMID were used (Table 3).

**Table 3.**
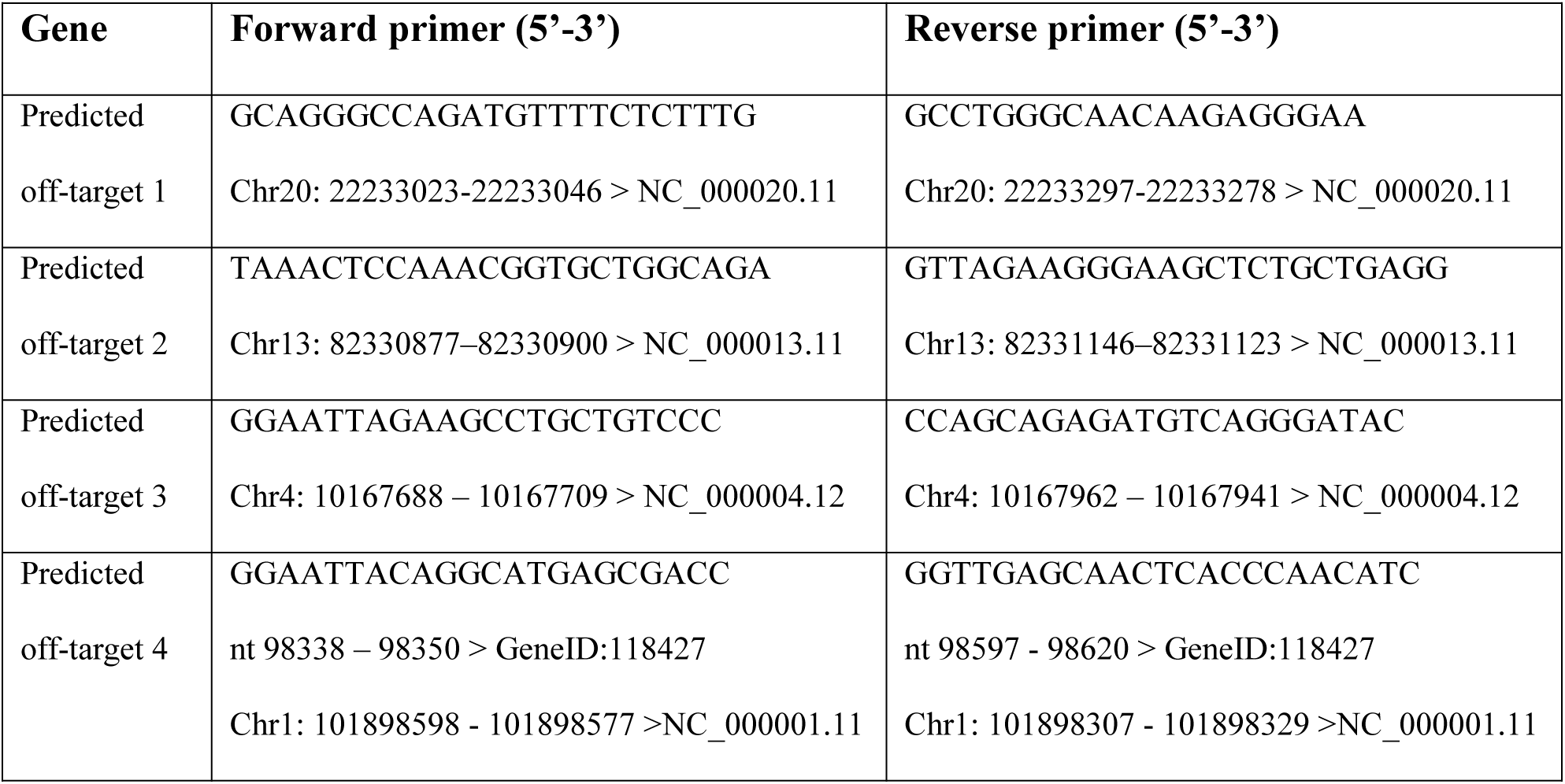
Primers used off-target sequencing of human RPTEC-TERT1 VPS33B-KO CRISPR clones.

### PCR clean-up & sequencing

For PCR clean up, MicroCLEAN (microzone, #2MCL-50) was used according to the manufacturer protocol. For off-target primers which generated multiple PCR products, so the QIAquick Gel Extraction Kit (QIAGEN, #28704) was used to extract the bands of interest from the gels according to the manufacturer protocol. Target genomic loci were then Sanger sequenced by Source Bioscience. Sequenced files were then analysed using the ICE Analysis tool from Synthego (16). To understand the result of mutations found during sequencing on protein translation, VectorBuilder’s DNA Translation Tool was used.

### RNA-seq of RPTEC-TERT1 gene edited cells

RPTEC-TERT1 Control, KO1, and KO2 clones were cultured to 70-80% confluency before RNA extraction with the RNeasy Plus Mini kit (QIAGEN, #74136). Four biological replicates were processed for sequencing for each clone. RNA integrity was assessed (RIN > 7.0) using the Qubit RNA BR assay alongside Agilent’s 4200 Tapestation. Library preparation was performed with the Watchmaker RNA Library Prep Kit with Polaris™ Depletion (product number BK0002-096). Sequencing was performed on the NextSeq 2000 platform from Illumina. Sequencing was conducted on the NextSeq 2000 instrument at a concentration of 800 pM, utilizing a 57-bps-end run with corresponding 8-bps unique dual sample indexes and unique molecular indexes (UMIs).

Initial bioinformatic analysis and quality control of FASTQ files was carried out using the nf-core/rnaseq pipeline v3.12.0 (17). Downstream analysis was carried out on R version 4.3.1. DESeq2 v1.42.1 was used to normalise read counts and carry out the differential gene expression analysis. For further quality control analyses, principal component analyses (PCA) and heatmaps were plotted and generated with pheatmap v1.0.12, DESeq2 (18). The default Wald test was used to compare the Control and Knockout samples. Approximate posterior estimation for generalised linear model (GLM) coefficients was used to shrink log-fold changes between samples. For gene ontology (GO) and Kyoto Encyclopedia of Genes and Genomes (KEGG) analysis, genes with an adjusted p-value of <0.05 and 1.2<|fold changes| were analysed using clusterProfiler v4.10.1 (19) and gprofiler2 v0.2.3 (20), respectively. Graphs and plots were generated with enrichplot v1.22.0 and ggplot2 v.3.5.1.

### RPTEC-TERT1 monolayer survival assay

A monolayer survival assay was conducted by seeding RPTEC-TERT1 cells in 96-well wells at 5x10^4^ cells/well in RPTEC-TERT1 media. Every day for 11 days of culture, the wells were imaged using an EVOS M5000 microscope at x4 and x10 magnification and monolayer ‘survival’ (intact monolayer) or ‘failure’ (peeling) was recorded.

### Immunofluorescence staining for adhesion complexes (Phospho-paxillin) in RPTEC cells

RPTEC-TERT1 cells were seeded at 5x10^4^ cells/well in a 96-well plate and cultured for 6 days. Cells were fixed using 4% PFA for 15 minutes at room temperature, washed three times with DPBS, and blocked with block solution (10% FBS, 1% BSA, 0.1% Tween20 in PBS) for 1 hour at room temperature. After washing the cells with 0.1% Triton-X in PBS, cells were incubated with 50 μL/well of rabbit anti-phospho-paxillin antibody (Abcam, #ab4832) at 1:100 at 4°C overnight. After washing, cells were incubated with 50 μL/well of Donkey anti-Rabbit Alexa Fluor™ Plus 647 secondary antibody (Thermo Fisher, #A32795; 1:200) and TRITC-conjugated Phalloidin (Merck, #90228; 1:1000) for 1 hour, followed by DAPI (Merck, #90229) staining for 5 minutes at room temperature.

Imaging was carried out using a Axio Observer 7 (Zeiss, #491917-0001-000) microscope at x40 magnification. Images were analysed with CellProfiler v.4.2.6. Phospho-paxillin-positive punctae were enhanced and isolated using EnhanceorSuppressFeatures and IdentifyPrimaryObjects functions. The area covered by the punctae and the number of punctae per image were automatically counted by CellProfiler. To calculate the number of phospho-paxillin-positive punctae per cell, the number of punctae was divided by the number of nuclei per image according to the DAPI channel. To calculate the mean phospho-paxillin-positive puncta size for each image, the total area covered by phospho-paxillin-positive punctae was divided by the number of punctae per image. To investigate, the association of phospho-paxillin-positive punctae with phalloidin staining, the phalloidin staining was dilated using the DilateImage function with a disk-shaped structuring element of size 12. Then, the co-localisation of the punctae with the dilated phalloidin stain was quantified per image.

### Cell detachment assay

A cell detachment assay was adapted from a previously published protocol (21). RPTEC-TERT1 cells were seeded at 1 x 10^5^ cells/well in a 96-well plate, with 11 wells seeded for each timepoint (0-10 minutes). After overnight culture, the wells were washed with DPBS and placed in an oven heated to 37 °C to maintain a stable temperature. 100 μL of 0.05% trypsin-EDTA pre-warmed to 37 °C was added to each well, except time-point 0, where DPBS was added. At 1-minute intervals, trypsin was removed and media added to stop trypsinization. Wells were then washed, fixed with 4% PFA for 10 minutes, and stained with 2.3% crystal violet for 15 minutes.

The plate was then washed under running water and left to dry on paper towels overnight. Then, 50 μL/well of 10% acetic acid was added to dissolve the crystal violet followed by 250 μL of distilled water. Absorbance was read at 570 nm on a plate reader (BioTek, Synergy HT). Cell adhesion was calculated as a percentage relative to the 0-minute timepoint, after subtracting background control absorbance (cell-free well treated with crystal violet).

### Cell adhesion assay

RPTEC-TERT1 cells were trypsinised and seeded at 1x10^5^ cells/well in a 96-well plate. These cells were allowed to attach overnight. Then, the cells were washed with DPBS three times before being fixed with 50 μL/well of 4% PFA at room temperature for 10 minutes. Then, crystal violet staining was carried out as described above.

### Statistical Analysis

All statistical analyses were carried out on GraphPad Prism Software v10.3.1. The normal distribution of the data was confirmed with a Shapiro-Wilk test. For normally distributed data, two-tailed t-tests or one-way ANOVAs with Tukey’s multiple comparisons test were used. For survival analyses a log-rank test was used. Quantitative data, displayed as bar and line charts, are expressed as means ± standard error of the mean (SEM; error bars). Data are reported to 2 significant figures.

## Results

### Generation of a Human Renal Proximal Tubule VPS33B-KO cell line

CRISPR-Cas9n double nickase technology was used to target mutations to the first exon of the *VPS33B* gene in RPTEC-TERT1 cells. Two *VPS33B* double nickase plasmid sets (h1 and h2), targeting an overlapping region at the start of exon one, were compared for their cutting efficiency using the T7EI assay. The h2 plasmids had a higher cutting efficiency than h1 as evidenced by the brighter 350 bp band and minimal wild-type *VPS33B* 650 bp band generated (Figure 1A). Thus, the following experiments used h2-transfected cells.

**Figure 1.**
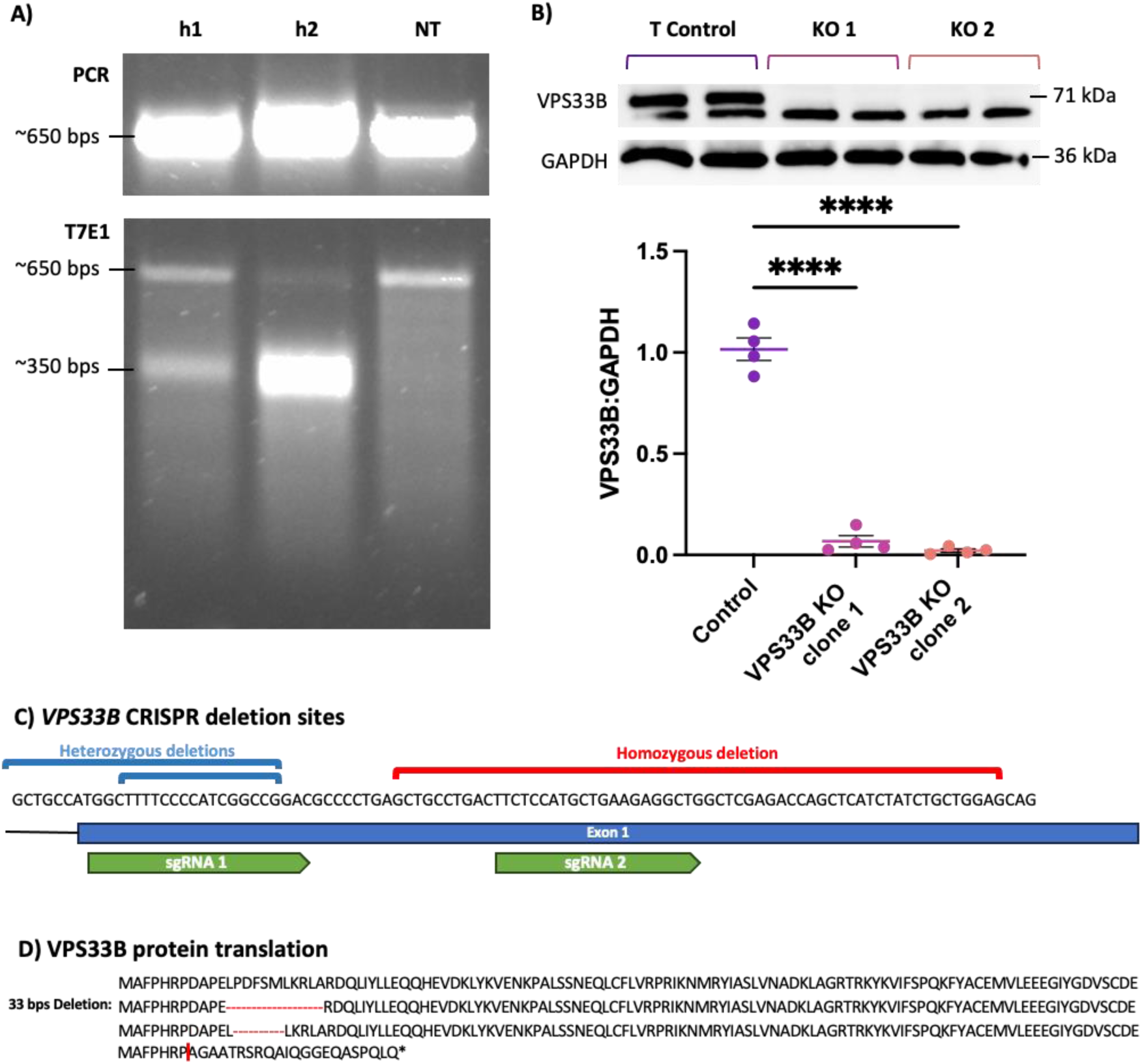
Generation and validation of VPS33B-knockout in a human renal proximal tubule epithelial cell (RPTEC) line. A) An image of the initial PCR reaction and subsequent T7EI assay carried out on gDNA collected from plasmid h1 transfected (left), plasmid h2 (middle), or untransfected (NT; right) RPTEC-TERT1 cells. B) Western blot (n=4) analysis of VPS33B expression in clones of a human proximal tubule cell line (RPTEC-TERT1). The RPTEC-TERT1 cells were either transfected with VPS33B-KO CRISPR plasmids resulting in successful knockout clones (VPS33B KO clones 1 & 2 – magenta & orange, respectively) or Control cells (purple). Western results were normalised against GAPDH expression. Data represents mean ± SEM. ****P<0.0001. ANOVA. Tukey’s multiple comparisons test. C) Sequence of 5’ of the first exon of the VPS33B gene and the deletions (compound heterozygotes – blue, homozygotes – red) identified in CRISPR-treated RPTEC-TERT1 cells. Exon 1 and the location of the CRISPR single-guide RNAs (sgRNAs) were depicted as a blue rectangle and green labels, respectively. D) Predicted changes to the amino acid sequence of VPS33B resulting from the mutations in B. Red dashes represent amino acid deletions and red line represents frameshift site.

Following CRISPR transfection, cells were fluorescence-activated cell sorted into 96-well plates to initiate clonal cell cultures. Clones were then expanded and genotyped. As ARC syndrome is an autosomal recessive disorder, a homozygous *VPS33B*-KO clone was desired. Sanger sequencing revealed that 2/8 selected clones were heterozygous, with insertions of either 3 bp or 14 bp in one allele. 3/8 clones were compound heterozygotes with the same 18 bps and 33 bps deletions (Figure 1C – blue). A further 2/8 clones were homozygous with 56 bp deletions in both alleles (Figure 1C – red), while a final clone contained no edits in *VPS33B* and is referred to from here on as the Control clone.

Vector Builder’s DNA translation tool predicted that the heterozygous 18 and 33 bps deletions would result in 6- or 11-amino acid deletions, respectively. Meanwhile, the 56-bps homozygous deletion was predicted to cause a frameshift resulting in a nonsense mutation and premature stop codon at amino acid 30 of the VPS33B sequence (Figure 1D). As the 56-bps homozygous deletion was predicted to essentially prevent protein translation, the two clones carrying this mutation were selected as the KO clones (KO clone 1 and KO clone 2).

Densitometry showed that VPS33B was significantly reduced in KO clone 1 (93% reduction, Dunnett’s test, p < 0.0001) and KO clone 2 (98% reduction, Dunnett’s test. p < 0.0001) compared to the Control (Figure 1B). Thus, the Western blotting demonstrated that the premature stop codon in the KO clone prevents VPS33B protein translation.

Finally, to explore the extent of off-target gene editing, likely off-target sites were predicted using the COSMID: CRISPR Search with Mismatches, Insertions and/or Deletions tool (14) and the WGE CRISPR Finder Tool. The top four predicted regions were Sanger sequenced, but no deletions or insertions were detected at these sites in either the Control line or the KO clones.

### Transcriptomic analysis of a VPS33B-KO RPTEC-TERT1 cell line

VPS33B is involved in a range of processes which differ from cell type to cell type and no specific information is available on its role in the proximal tubule. Thus, transcriptomic analysis of the RPTEC-TERT1 clones was used to identify potential mechanisms which may underlie the proximal tubular dysfunction seen in ARC syndrome.

Principal component analysis (PCA) demonstrated clear clustering of the knockout clones along PC1, accounting for 65% of the variability in the data (Figure 2A). Hierarchical clustering was also carried out to generate pairwise comparisons of the expression profiles of the samples. This also demonstrated that the KO clones clustered together, separately from the control (Figure 2C). Furthermore, correlation coefficients observed in this hierarchical clustering analysis showed high correlation (>0.998) in all pairwise comparisons, suggesting no outlying samples (Figure 2B). To reduce noise and increase the power of the differential gene expression analysis, subsequent analyses were carried out comparing the control line against the two *VPS33B*-KO clones pooled. Analysis of differential gene expression showed a high number of differentially expressed genes (DEGs) with relatively low fold-changes. Thresholds of adjusted p < 0.05 and |fold change| > 1.2 were selected. With this fold-change threshold, a total of 3,362 genes were found to be differentially expressed with 1,790 being upregulated and 1,572 being downregulated in *VPS33B*-KO samples compared to the Control samples (Figure 2C).

**Figure 2.**
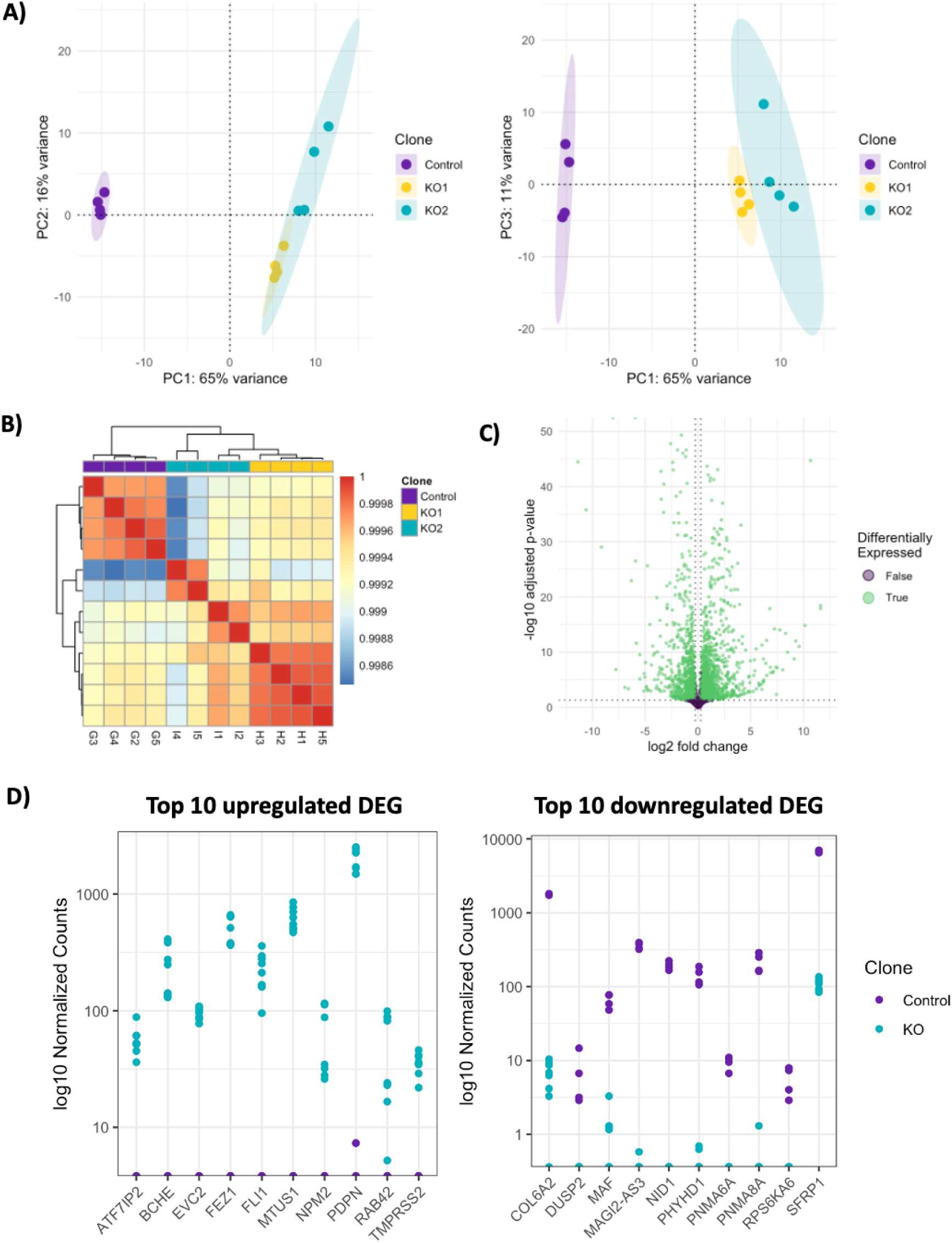
Exploratory transcriptomic analyses of control and VPS33B-knockout RPTEC-TERT1 cells. A) Principal component analysis control and knockout transcriptomes. Top principal components totalled 92% and the x axis in both PCA plots (PCA1) and the y axis in the left PCA plot (PCA2) and right PCA (PCA3) plot, respectively, as follows: 1=65%, 2=16%, and 3=11%. B) A heatmap with a dendrogram demonstrating the similarity of the control samples (G2-5) to the two knockout clones (KO1: H1-5 and KO2:I1-5). Colouring denotes correlation coefficient (red: high, blue: low). C) A volcano plot of differentially expressed genes between control and knockout clones (P < 0.05, |Fold-change| > 1.2). False: purple, True: green. D) Plot of log10 normalised counts of top 10 up- and down-regulated (by fold-change) differentially expressed genes. Abbreviations: DEG: Differentially expressed genes.

The top 10 upregulated DEGs, ranked by largest fold change were: *ATF7IP2* (log2FC = 9.27), *BCHE* (log2FC = 11.6), *EVC2* (log2FC = 10.1), *FEZ1* (log2FC = 12.8), *FLI1* (log2FC = 11.6), *MTUS1* (log2FC = 13.2), *NPM2* (log2FC = 9.59), *RAB42* (log2FC = 9.11), *PDPN* (log2FC = 10.7), and *TMPRSS2* (log2FC = 8.50) (Figure 2E). The top 10 downregulated DEGs were: *COL6A2* (log2FC = -8.06), *DUSP2* (log2FC = -6.93), *MAF* (log2FC = -6.32), *MAGI2-AS3* (log2FC = -11.4), *NID1* (log2FC = -13.0), *PHYHD1* (log2FC = -9.16), *PNMA6A* (log2FC = 7.78), *PNMA8A* (log2FC = -10.6), *RPS6KA6* (log2FC = -6.60), and *SFRP1* (log2FC = -5.94) (Figure 2D).

To understand the functional roles of the DEGs in the *VPS33B*-KO cells, gene ontology (GO) and Kyoto Encyclopedia of Genes and Genomes (KEGG) analyses were carried out to investigate the DEGs’ association to biological processes and identify enriched pathways, respectively. GO and KEGG analyses each retrieved 503 and 273 significantly enriched terms (adjusted p-value < 0.05). To investigate the most significantly enriched terms, the top 20 (by adjusted p-value) GO and KEGG terms were investigated.

Both the top 20 KEGG and top 20 GO terms showed enrichment of cell adhesion-associated pathways (GO: “positive regulation of cell adhesion”; KEGG: “Focal adhesion” and “ECM-receptor interaction”). The top 20 GO terms also included 11 terms associated with kidney development or development in general (“kidney development”, “renal system development”, “renal tubule development”, “nephron development”, “renal epithelium development”, “nephron tubule development”, “embryonic organ development”, “renal tubule morphogenesis”, “nephron epithelium development”, “kidney morphogenesis”, and “epithelial tube morphogenesis”) and two terms associated with ossification (“ossification” and “osteoblast differentiation” (Figure 3A,B). Meanwhile, in addition to the cell adhesion-associated terms, the KEGG analysis showed enrichment of cancer associated pathways (“Pathways in cancer”, “MAPK signaling pathway”, “PI3K-Akt signaling pathway”, “Human papillomavirus infection”, “Proteoglycans in cancer”, “Rap1 signaling pathway”, “Hippo signaling pathway”, “Transcriptional misregulation in cancer”, “Hepatocellular carcinoma”, “Ras signaling pathway”, “Gastric cancer”), and infection associated pathways (“Efferocytosis”, “Salmonella infection”) in the top 20 KEGG terms (Figure 3C).

**Figure 3.**
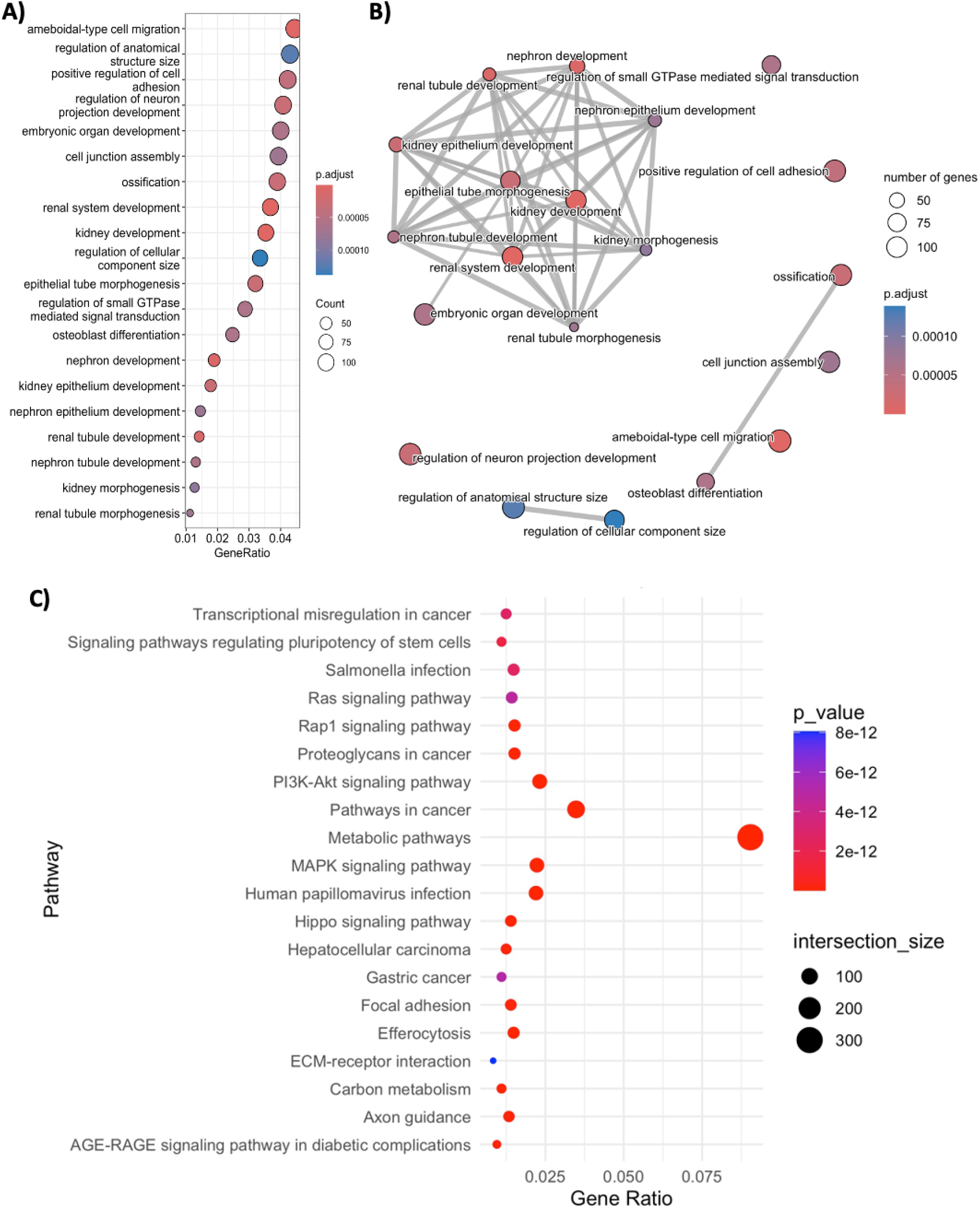
Pathway analysis of VPS33B-knockout RPTEC-TERT1 cells. A) Dot plot of the top 20 enriched gene ontology terms (by adjusted p-value) comparing control to knockout clones. B) Enrichment map of top 20 gene ontology terms. Lines (edges) between nodes represent pairwise Jaccard coefficients, with a default threshold of 0.2 for drawing connections. The Jaccard coefficient is calculated as the ratio of the total number of genes shared between terms to the total number of unique genes associated with both terms. Thicker edges indicate higher Jaccard coefficients, while thinner edges indicate lower coefficients. Dot plot of 20 C) enriched KEGG terms (by adjusted p-value). For dotplots A and C: Gene ratio: Number of DEG matching a term/total number of DEG. Graphs were coloured according to their adjusted p-values with blue denoting higher p-values and red denoting lower p-values. Intersection size (size of nodes) refers to the number of DEG matching a term. Abbreviations: DEG: Differentially expressed genes.

### General phenotypic characterisation of VPS33B-KO RPTEC-TERT1 cell line in extended culture

Next, for an initial phenotypic characterisation of *VPS33B*-KO RPTEC-TERT1 clones, RPTEC-TERT1 cells were cultured in a 96-well plate for 11 days and observed daily with brightfield microscopy. At 1 day following seeding, all clones appeared elongated (blue arrows in Figure 4). By day 3 of culture, cells reached confluency and by day 5, they reached the typical epithelial cobblestone shape expected of RPTEC-TERT1 cells (22). No obvious differences in cell behaviour were noted between Control and KO clones until day 6 of culture, when *VPS33B*-KO but not Control cells were seen to progressively begin to ‘peel’ as a monolayer from the tissue culture plastic (black arrows, Figure 4B,C).

**Figure 4.**
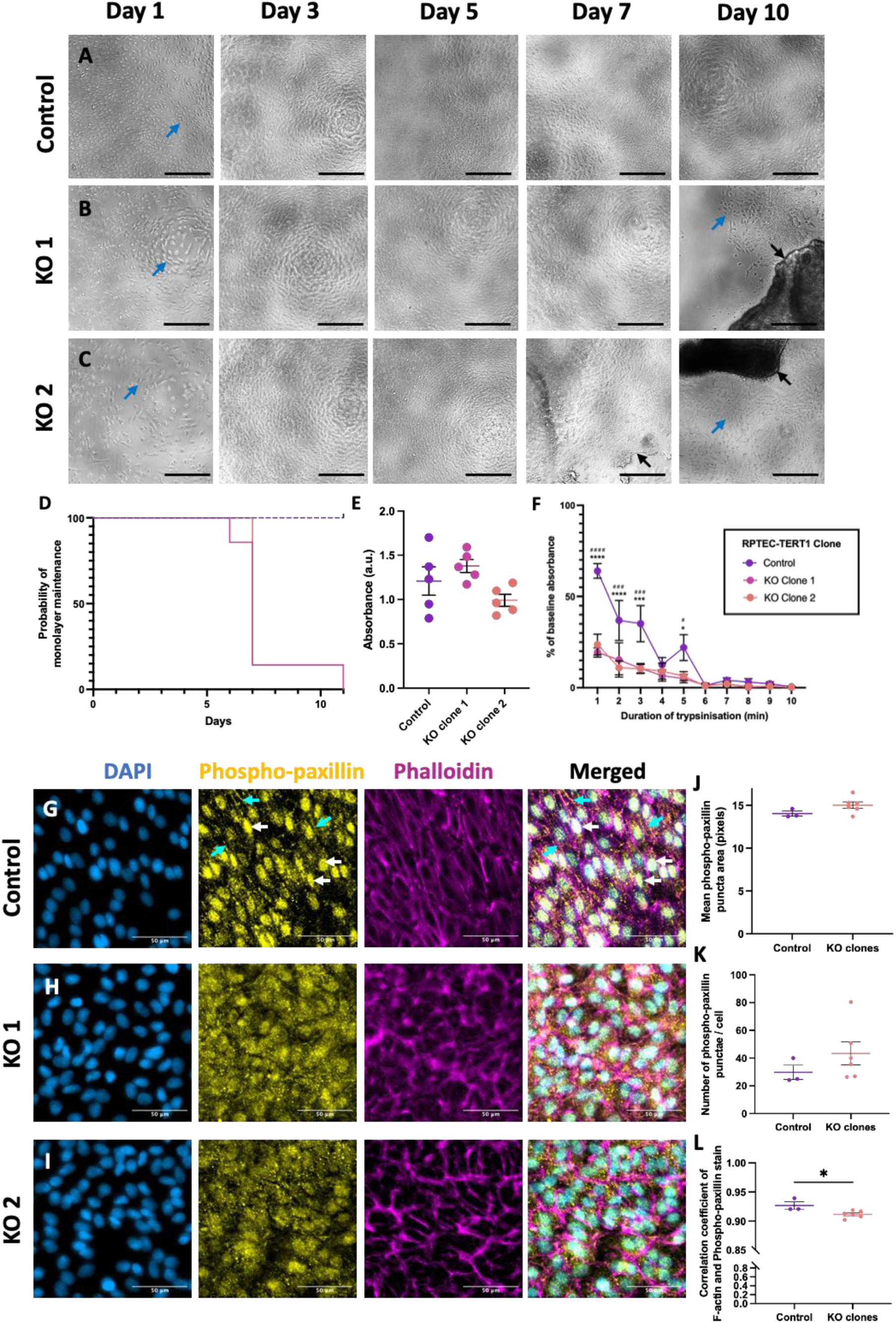
VPS33B-KO RPTEC-TERT1 clones demonstrate weaker attachment compared to controls. A) x10 brightfield images of A) Control, B) VPS33B-KO clone 1, and C) VPS33B-KO clone 2 RPTEC-TERT1 cells in 96-well plates after 1, 3, 5, 7, and 10 days of culture following seeding at 5x10^4^/well. Blue arrows: elongated RPTEC-TERT1 cells. Black arrow: Monolayer peeling. Scale bar: 100 µm. D) Survival analysis for time until peeling for clones of a human proximal tubule cell line (RPTEC-TERT1). The RPTEC-TERT1 cells were either transfected with VPS33B-KO CRISPR plasmid resulting in successful knockout clones (VPS33B KO clones 1 & 2 – magenta & orange, respectively) or wild-type but transfected cells (Control - purple). n=7. Log-rank test for trend. p < 0.0001. E) Analysis of absorbance of crystal violet one day after seeding 1x10^5^ cells/well. ANOVA. Tukey’s multiple comparisons test. n=5. F) Results of a cell detachment assay carried out on Control and VPS33B-KO RPTEC-TERT1 cells. Crystal violet absorbance is used to measure the number of cells attached after 1-10 minutes of trypsinisation. Two-way ANOVA. Tukey’s multiple comparisons test. Control vs KO clone 1: ^#^p<0.05; ^###^p<0.001, ^####^p<0.0001. Control vs KO clone 2: *p<0.05; ***p<0.001, ****p<0.0001. n = 5. Representative x40 images of RPTEC-TERT1 cells stained with DAPI (blue), anti-phospho-paxillin antibody (yellow), and phalloidin-stained F-actin (magenta) at 5 days after seeding. Cyan arrows indicate linear junctional staining of phospho-paxillin. White arrows indicate nuclear phospho-paxillin staining. G) Control: CRISPR-transfected control, H) KO1: VPS33B-KO RPTEC-TERT1 Clone 1, I) KO2: VPS33B-KO RPTEC-TERT1 Clone 2. n=3. Scale bar = 50 µm. Analysis of J) mean size of phospho-paxillin-positive puncta area, K) number of punctae per cell, and L) association of phospho-paxillin-positive punctae with the actin cytoskeleton comparing Control cells to a pooled sample of VPS33B-KO clones. n=3 per genotype. T-test. Data represents mean ± SEM. *p<0.05. Abbreviations: a.u.: arbitrary units.

To quantify this peeling phenotype, RPTEC-TERT1 cells were cultured for up to 11 days and the number of days taken for each clone to peel from the tissue culture plastic was counted for 7 independent replicates. This was used to calculate the probability of monolayer maintenance over time with a peeling event constituting monolayer “failure”. While the Control line did not peel during the 11-day period studied, both of the *VPS33B*-KO lines peeled completely within 11 days (Log-rank test for trend, p < 0.0001; Figure 4D). Most *VPS33B*-KO cells peeled at 7 days post seeding while some wells remained unpeeled for up to 10 days. Brightfield imaging showed that RPTEC-TERT1 cells appeared elongated initially upon seeding and became smaller and more regular after reaching confluency around day 3 of culture. Both *VPS33B*-KO clones began peeling around day 7 after which elongated cells invaded the space left behind by the peeled monolayer (blue arrows, Figure 4B,C).

### Cell attachment assays

To investigate the mechanobiological mechanism underlying the monolayer peeling observed in the KO clones, cell attachment and cell detachment assays were carried out. To investigate whether the *VPS33B* KO reduced the ability of cells to initially adhere to their substrate upon seeding, cells were seeded into 96-well plates. The following day, they were washed with DPBS to remove any cells which had not attached and stained with crystal violet (Figure 4E). No significant differences in cells’ ability to attach to the tissue culture surface were observed between the Control and KO clones (ANOVA. p>0.05). To investigate this further, analysis of the strength of cell adhesion was carried out, using a cell detachment assay. Cells were seeded into a 96-well plate and allowed to attach overnight. These cells were then trypsinised for 0, 1, 2, 3, 4, 5, 6, 7, 8, 9, or 10 minutes and a crystal violet stain was used to visualise the cell density remaining (Figure 4F).

Results showed that both *VPS33B*-KO clones took significantly less time (both reaching <20% attachment in 1 minute) to detach compared to the transfected control (reaching <20% attachment after 6 minutes; (Figure 4F). There was no significant difference in the rate of cell detachment between the two VPS33B-KO clones at any time point, with KO clone 1 and KO clone 2 being 81% and 76% detached within the first minute of trypsinisation compared to 36% in the Control line. At minute 1 of trypsinisation a Tukey’s multiple comparisons test showed significant differences in cell detachment between Control and KO clone 1 (p < 0.0001), and Control and KO clone 2 (p < 0.0001). This significant difference in the rate of trypsinisation could be observed until minute 5 of trypsinisation. Unlike the cell attachment assay, this demonstrated a clear demarcation between both Control and KO clones, suggesting that *VPS33B* KO results in the strength of cell-substrate adhesion.

### Adhesion complex distribution is disrupted in VPS33B-KO RPTEC-TERT1 cells

To better understand the source of this attachment defect, we investigated focal adhesions in VPS33B-KO RPTEC-TERT1 cells. Knockdown of focal adhesion components, such as integrin β3 in Chinese Hamster Ovary cells, can reduce the ability of cells to attach to substrates (23). Thus, it was hypothesized that the adhesion defect would be associated with a change in focal adhesion component distribution.

Immunofluorescent staining with anti-phospho-paxillin antibodies and phalloidin visualised focal adhesions and actin, respectively. Control cells showed nuclear (white arrows) and cortical (cyan arrows) localisation of phospho-paxillin with punctate staining associating with the actin cytoskeleton (Figure 4G). Meanwhile, both KO lines showed more diffuse, cytoplasmic, punctate staining of phospho-paxillin with no obvious cortical localisation (Figure 4H,I).

To investigate whether an automated image analysis software could differentiate between Control and KO clones, CellProfiler software was used to analyse the images and the results of the Control and pooled KO clones were compared. Analysis found no significant differences in number (Figure 4J) or size (Figure 4K) of phospho-paxillin punctae between Control and KO clones. To quantify the association of the focal adhesion with the actin cytoskeleton, the co-localisation of the phospho-paxillin and phalloidin stain were quantified. This analysis demonstrated that *VPS33B* KO significantly reduced phospho-paxillin proximity to the actin cytoskeleton (Figure 4L; p = 0.027) without reducing the number or size of the phospho-paxillin punctae.

## Discussion

This study generated *VPS33B*-KO RPTEC-TERT1 cells which were found to have an adhesion defect. Both KEGG and GO analyses of these cells demonstrated an enrichment of cell adhesion-associated terms, and spontaneous monolayer peeling and cell adhesion assays provided functional evidence of a cell adhesion phenotype.

The KEGG and GO analyses showed similar trends to other *VPS33B* depletion models. GO analysis of differentially expressed genes uncovered changes associated with renal and epithelial development/tube formation, regulation of cell and anatomical structure size, ameboidal-type cell migration, ossification, neuronal development and cell adhesion. Meanwhile, KEGG analysis showed enrichment in ECM-receptor interactions and focal adhesions, infection-associated (HIV and HPV infections) and cancer-associated pathways. Similar pathways were found to be dysregulated in Chai et al. who examined *Vps33b^fl/fl^; Alfp-Cre (hepatocyte-KO)* mouse livers (24,25), with their top 5 GO terms being “Cell adhesion”, “Biological adhesion”, “Defence response”, “Response to wounding”, and “Immune response”. Their top 5 enriched pathways were “ABC transporters” “Cell adhesion”, “ECM receptor interaction”, “Toll-like receptor signalling pathway”, and “Focal adhesion” (24). Despite the differences in species and organ, there is commonality in the enriched adhesion and inflammatory/immune GO and KEGG terms seen in Figure 3. Proteomic comparison of *Vps33b*-KO and *Vps33b*-overexpressing immortalised mouse tendon fibroblasts also showed enrichment of adhesion-related GO terms (26). The similarities of the results of the RNA-seq analysis carried out in this paper with previous omic studies of *Vps33b*-KO cells and tissues supports the notion that the analysis successfully identified key VPS33B-regulated processes.

There is little previous literature on the role of VPS33B in cell-substrate adhesion. However, it is known that VPS33B can bind to integrins β1 and β3, which as part of integrins α_v_β_3_ and α_5_β_1_, are found in focal adhesions (27,28). It has also been shown in murine cells that Vps33b is involved in integrin αIIbβ3-mediated endocytosis and integrin outside-in signalling, which affects cell spreading (27). It is not entirely clear what the role of VPS33B in integrin outside-in signalling is. It is unlikely to be as simple as trafficking the integrins, as similar levels of activated integrin αIIbβ3 were found on control and *Vps33b*-KO mouse platelets in response to thrombin stimulation, meaning cells did not have problems trafficking this integrin to their cell surfaces (27). However, our paper provides the first evidence of a direct role for VPS33B in cell adhesion.

Phospho-paxillin staining showed changes in response to *VPS33B* KO. Phospho-paxillin staining was focal adhesion-like (punctate), associating with the actin cytoskeleton in the control clones. However, the pattern was consistently disrupted in *VPS33B*-KO RPTEC-TERT1 cells. This suggests that focal adhesion protein distribution is disrupted by *VPS33B* KO. Additionally, the lack of a significant difference in the number of phospho-paxillin-positive punctae between *VPS33B*-KO and Control cells suggests that *VPS33B* KO does not affect focal adhesion number, only their distribution to points of adhesion.

This adhesion defect may provide the first insights into the pathophysiology of ARC syndrome in the kidney. The proximal tubule receives the largest volume of fluid of all the nephron components, being the first section of the nephron after the Bowman’s capsule (29). Therefore, it is likely to experience the highest shear stress in the nephron. Taken together, we propose a model in which VPS33B loss causes an adhesion defect in proximal tubule epithelial cells, rendering them more vulnerable to shear stress. This shear stress may lead to persistent tissue injury and inflammation *in vivo* and ultimately cause the tubular damage seen in ARC syndrome. To confirm the role of VPS33B-related adhesion defects in renal ARC syndrome pathogenesis, future studies could examine focal adhesion organisation and cellular responses to shear stress in patient samples. Haemorrhages seen in ARC syndrome patients prevent the collection of biopsy samples, but such research could be carried out on autopsy samples or cells captured from patients’ urine (30).

## Conclusion

This study demonstrates a critical role for VPS33B in cell adhesion within renal proximal tubule cells, providing novel insights into the pathophysiology of ARC syndrome. Characterisation of *VPS33B*-KO RPTEC-TERT1 cells revealed significant cell adhesion defects, supported by RNA-seq analysis, detachment assays, and disrupted phospho-paxillin staining patterns. These findings suggest that VPS33B deficiency compromises cell-matrix interactions, potentially rendering proximal tubule cells more susceptible to damage *in vivo* and contributing to the renal dysfunction observed in ARC syndrome.

## CRediT authorship contribution statement

**Maria Caluianu:** Conceptualization, Investigation, Methodology, Visualisation, Writing – original draft, Writing – review & editing. **Kimberley A. Owen:** Visualisation, Writing – review & editing.

## Data Availability

The RNA-seq datasets generated during this study will be made publicly available in the Gene Expression Omnibus (GEO) repository upon publication. The GEO accession number will be provided in the final published article

## Acknowledgements

Thank you to Dr. Robert E. Hynds (University College London, UK) for providing feedback on this paper and for the support he provided during the publication period. We thank UCL Genomics for their support with the sequencing in this study.

## Supplementary data

## References

1. Gissen P, Tee L, Johnson CA, Genin E, Caliebe A, Chitayat D, et al. Clinical and molecular genetic features of ARC syndrome. Hum Genet. 2006 Oct;120(3):396–409.

2. Cullinane AR, Straatman-Iwanowska A, Zaucker A, Wakabayashi Y, Bruce CK, Luo G, et al. Mutations in VIPAR cause an arthrogryposis, renal dysfunction and cholestasis syndrome phenotype with defects in epithelial polarization. Nat Genet. 2010 Apr;42(4):303–12.

3. Zhou Y, Zhang J. Arthrogryposis-renal dysfunction-cholestasis (ARC) syndrome: from molecular genetics to clinical features. Ital J Pediatr. 2014 Sep 20;40:77.

4. Smith H, Galmes R, Gogolina E, Straatman-Iwanowska A, Reay K, Banushi B, et al. Associations among genotype, clinical phenotype, and intracellular localization of trafficking proteins in ARC syndrome. Hum Mutat. 2012 Dec;33(12):1656–64.

5. Tornieri K, Zlatic SA, Mullin AP, Werner E, Harrison R, L’hernault SW, et al. Vps33b pathogenic mutations preferentially affect VIPAS39/SPE-39-positive endosomes. Hum Mol Genet. 2013 Dec 20;22(25):5215–28.

6. Lo B, Li L, Gissen P, Christensen H, McKiernan PJ, Ye C, et al. Requirement of VPS33B, a member of the Sec1/Munc18 protein family, in megakaryocyte and platelet alpha-granule biogenesis. Blood. 2005 Dec 15;106(13):4159–66.

7. Satomura Y, Bessho K, Nawa N, Kondo H, Ito S, Togawa T, et al. Novel gene mutations in three Japanese patients with ARC syndrome associated mild phenotypes: a case series. J Med Case Reports. 2022 Feb 13;16(1):60.

8. Penon-Portmann M, Westbury SK, Li L, Pluthero FG, Liu RJY, Yao HHY, et al. Platelet VPS16B is dependent on VPS33B expression, as determined in two siblings with arthrogryposis, renal dysfunction, and cholestasis syndrome. J Thromb Haemost. 2022 Mar 24;

9. Dai J, Lu Y, Wang C, Chen X, Fan X, Gu H, et al. Vps33b regulates Vwf-positive vesicular trafficking in megakaryocytes. J Pathol. 2016 Sep;240(1):108–19.

10. Smith H. INVESTIGATION OF VPS33B DEFICIENCY IN MOUSE AND MAN [Doctoral dissertation]. The University of Birmingham; 2012.

11. Banushi B, Forneris F, Straatman-Iwanowska A, Strange A, Lyne A-M, Rogerson C, et al. Regulation of post-Golgi LH3 trafficking is essential for collagen homeostasis. Nat Commun. 2016 Jul 20;7:12111.

12. Wieser M, Stadler G, Jennings P, Streubel B, Pfaller W, Ambros P, et al. hTERT alone immortalizes epithelial cells of renal proximal tubules without changing their functional characteristics. Am J Physiol Renal Physiol. 2008 Nov;295(5):F1365–75.

13. Perretta-Tejedor N, Freke G, Seda M, Long DA, Jenkins D. Generating mutant renal cell lines using CRISPR technologies. Methods Mol Biol. 2020;2067:323–40.

14. Cradick TJ, Qiu P, Lee CM, Fine EJ, Bao G. COSMID: A Web-based Tool for Identifying and Validating CRISPR/Cas Off-target Sites. Mol Ther Nucleic Acids. 2014 Dec 2;3(12):e214.

15. Hodgkins A, Farne A, Perera S, Grego T, Parry-Smith DJ, Skarnes WC, et al. WGE: a CRISPR database for genome engineering. Bioinformatics. 2015 Sep 15;31(18):3078– 80.

16. Conant D, Hsiau T, Rossi N, Oki J, Maures T, Waite K, et al. Inference of CRISPR Edits from Sanger Trace Data. The CRISPR Journal. 2022 Feb 2;5(1):123–30.

17. Ewels PA, Peltzer A, Fillinger S, Patel H, Alneberg J, Wilm A, et al. The nf-core framework for community-curated bioinformatics pipelines. Nat Biotechnol. 2020 Mar;38(3):276–8.

18. Love MI, Huber W, Anders S. Moderated estimation of fold change and dispersion for RNA-seq data with DESeq2. Genome Biol. 2014;15(12):550.

19. Wu T, Hu E, Xu S, Chen M, Guo P, Dai Z, et al. clusterProfiler 4.0: A universal enrichment tool for interpreting omics data. Innovation (Camb). 2021 Aug 28;2(3):100141.

20. Kolberg L, Raudvere U, Kuzmin I, Vilo J, Peterson H. gprofiler2 -- an R package for gene list functional enrichment analysis and namespace conversion toolset g:Profiler. F1000Res. 2020 Jul 15;9.

21. Lau CH, Rouhani MJ, Maughan EF, Orr JC, Kolluri KK, Pearce DR, et al. Lentiviral expression of wild-type LAMA3A restores cell adhesion in airway basal cells from children with epidermolysis bullosa. Mol Ther. 2024 May 1;32(5):1497–509.

22. Aschauer L, Gruber LN, Pfaller W, Limonciel A, Athersuch TJ, Cavill R, et al. Delineation of the key aspects in the regulation of epithelial monolayer formation. Mol Cell Biol. 2013 Jul;33(13):2535–50.

23. Bhattacharya R, Gonzalez AM, Debiase PJ, Trejo HE, Goldman RD, Flitney FW, et al. Recruitment of vimentin to the cell surface by beta3 integrin and plectin mediates adhesion strength. J Cell Sci. 2009 May 1;122(Pt 9):1390–400.

24. Chai M, Su L, Hao X, Zhang M, Zheng L, Bi J, et al. Identification of genes and signaling pathways associated with arthrogryposis-renal dysfunction-cholestasis syndrome using weighted correlation network analysis. Int J Mol Med. 2018 Oct;42(4):2238–46.

25. Hanley J, Dhar DK, Mazzacuva F, Fiadeiro R, Burden JJ, Lyne A-M, et al. Vps33b is crucial for structural and functional hepatocyte polarity. J Hepatol. 2017 May;66(5):1001–11.

26. Chang J, Pickard A, Herrera JA, O’Keefe S, Hartshorn M, Garva R, et al. Endocytic recycling is central to circadian collagen fibrillogenesis and disrupted in fibrosis. 2024 Apr 3;

27. Xiang B, Zhang G, Ye S, Zhang R, Huang C, Liu J, et al. Characterization of a novel integrin binding protein, VPS33B, which is important for platelet activation and in vivo thrombosis and hemostasis. Circulation. 2015 Dec 15;132(24):2334–44.

28. Rossier O, Octeau V, Sibarita J-B, Leduc C, Tessier B, Nair D, et al. Integrins β1 and β3 exhibit distinct dynamic nanoscale organizations inside focal adhesions. Nat Cell Biol. 2012 Oct;14(10):1057–67.

29. Gilmer GG, Deshpande VG, Chou C-L, Knepper M. Flow resistance along the rat renal tubule. Am J Physiol Renal Physiol. 2018 Nov 1;315(5):F1398–405.

30. Nazmutdinova K, Man CY, Carter M, Beales PL, Winyard PJ, Walsh SB, et al. Cell Catcher: a new method to extract and preserve live renal cells from urine. medRxiv. 2023 Sep 22;

